# Spatial organization of face part representations within face-selective areas revealed by high-field fMRI

**DOI:** 10.1101/2021.06.23.449598

**Authors:** Jiedong Zhang, Yong Jiang, Yunjie Song, Peng Zhang, Sheng He

**Affiliations:** Institute of Biophysics, Chinese Academy of Sciences, Beijing 100101, China; University of Chinese Academy of Sciences, Beijing 100049, China; Department of Psychology, University of Minnesota, Minneapolis, MN 55455

**Keywords:** face processing, high-field fMRI, spatial organization, FFA

## Abstract

Regions sensitive to specific object categories as well as organized spatial patterns sensitive to different features have been found across the whole ventral temporal cortex (VTC). However, it is unclear that within each object category region, how specific feature representations are organized to support object identification. Would object features, such as object parts, be represented in fine-scale spatial organization within object category-specific regions? Here we used high-field 7T fMRI to examine the spatial organization of neural tuning to different face parts within each face-selective region. Our results show consistent spatial organization across individuals that within right posterior fusiform face area (pFFA) and right occipital face area (OFA), the posterior portion of each region was biased to eyes, while the anterior portion was biased to mouth and chin stimuli. Our results demonstrate that within the occipital and fusiform face processing regions, there exist systematic spatial organizations of neural tuning to different face parts that support further computation combining them.

## Introduction

The ventral temporal cortex (VTC) in the brain supports our remarkable ability to recognize objects rapidly and accurately from the visual input in everyday life. Identity information is extracted from visual input through multiple stages of representation. To fully understand the neural mechanism of object processing, it is critical to know how these representations are physically applied to anatomical neural structure in the VTC. Numerous studies have already revealed multiple levels of feature representation manifest at different scales of anatomical organizations which superimposed in the VTC. The superordinate category representations (e.g., animate/inanimate, real-world size) manifest at large scale organization covering the whole VTC. Meanwhile, the category-selective representations (e.g., face, body, and scene selective regions in the mid-fusiform gyrus) are revealed at finer spatial scale in the VTC (1, 2). Recent evidence suggested a general spatial organization of neural responses to dimensions in object feature space in monkey inferotemporal cortex (3). Could such physical organization be further extended to even smaller scale, like object parts/features representations within each category-selective region? In other words, as part representations play a critical role in object processing, would there be consistent spatial organizations across individuals for different object parts within each category-selective region in VTC?

Fine-scale spatial organizations of low-level visual features have already been found in early visual cortex, such as ocular dominance columns and orientation pinwheels (4–7). Among all the object-selective regions in the VTC, the face-selective regions, including FFA and OFA, are one of the most widely examined object-processing networks in the past decades in cognitive neuroscience. As faces have spatially separated yet organized features such as eyes and mouth which are easy to be defined, it is suitable to use face parts to examine whether there are spatial organizations for different object features in the VTC. Neurophysiology studies in non-human primates demonstrated face-selective neurons in face-selective regions showed different sensitivities to various of face feature or combination of dimensions in face feature space (8, 9). Human fMRI studies also found the neural response patterns in FFA or OFA could distinguish different face parts (10), suggesting voxels within same face-selective region may have different face feature tuning. In addition, previous study also suggests that the spatial distribution of a face feature may be relevant to the physical location of that feature in a face (11).

The sizes of the face-selective regions in VTC are relatively small, spanning about 1 cm. To investigate the potential spatial organization within each face region, high-resolution fMRI with sufficient sensitivity and spatial precision is necessary. With high-field fMRI, fine-scale patterns have been observed in early visual cortex, such as columnar-like structures in V1, V2, V3, V3a, and hMT (12–17). These findings validate the feasibility of using high-field fMRI to reveal fine-scale (several mm) structures in the visual cortex.

Here we used 7T fMRI to examine whether category-specific feature information, such as object parts, would be represented in certain spatial organization within object selective regions. With faces as stimuli, the high-field fMRI allowed for measuring detailed neural response patterns from multiple face-selective regions. Our results show that in the right pFFA and right OFA, different face parts elicited differential spatial patterns of fMRI responses. Specifically, eyes induced responses biased to the posterior portion of the ROIs while responses to mouth and chin were biased to the anterior portion of the ROIs. Similar face parts based spatial organizations were observed in both the pFFA and OFA, and the patterns are highly consistent across participants. Together, these results reveal robust fine-scale spatial organizations of face feature representation within face-selective regions.

## Results

One critical challenge to demonstrate the spatial organization within single face-selective region is to find the anatomical landmark to align the function maps between different individuals, as the shape, size, and spatial location of FFA vary largely across individuals. Among all the anatomical structures in the VTC, the mid-fusiform sulcus (MFS) could potentially serve as landmark in the current study. MFS is relatively small structure in the VTC, but consistently present in most individuals (18). On the one hand, the structure of MFS could predict the coordinates of face-selective region around mid-fusiform, especially the anterior one (18). On the other hand, MFS is found to be highly consistent with many anatomical lateral-medial transitions in the VTC, such as cytoarchitecture and white-matter connectivity transitions (18–21). In addition, it could also predict the transitions in many function organization, such as animacy/inanimacy and face/scene preference (20). Considering its anatomical and functional significance, in the current study, we used the direction of MFS to align the potential spatial organization of face part across individuals.

Different face parts (i.e., eyes, nose, mouth, hair, and chin, see Figure 1A) and whole faces were presented to participants and they performed a one-back task in 7T MRI scanner. For each participant, five face-selective ROIs (i.e., right pFFA, right aFFA, right OFA, left FFA, and left OFA) were defined with independent localizer scans. Before comparing the spatial response patterns between the face parts, we assessed the overall neural response amplitudes they generated in each ROIs. All face selective regions showed a similar trend that eyes generated higher responses than nose, hair, and chin (ts>2.61, ps<0.05; except for eyes vs. nose in the left FFA and for eyes vs. chin in left OFA, ts<2.40, ps> 0.06. See Figure 1B). However, mouth generated similar response amplitudes as eyes (ts<1.58, ps>0.17).

**Figure 1.**
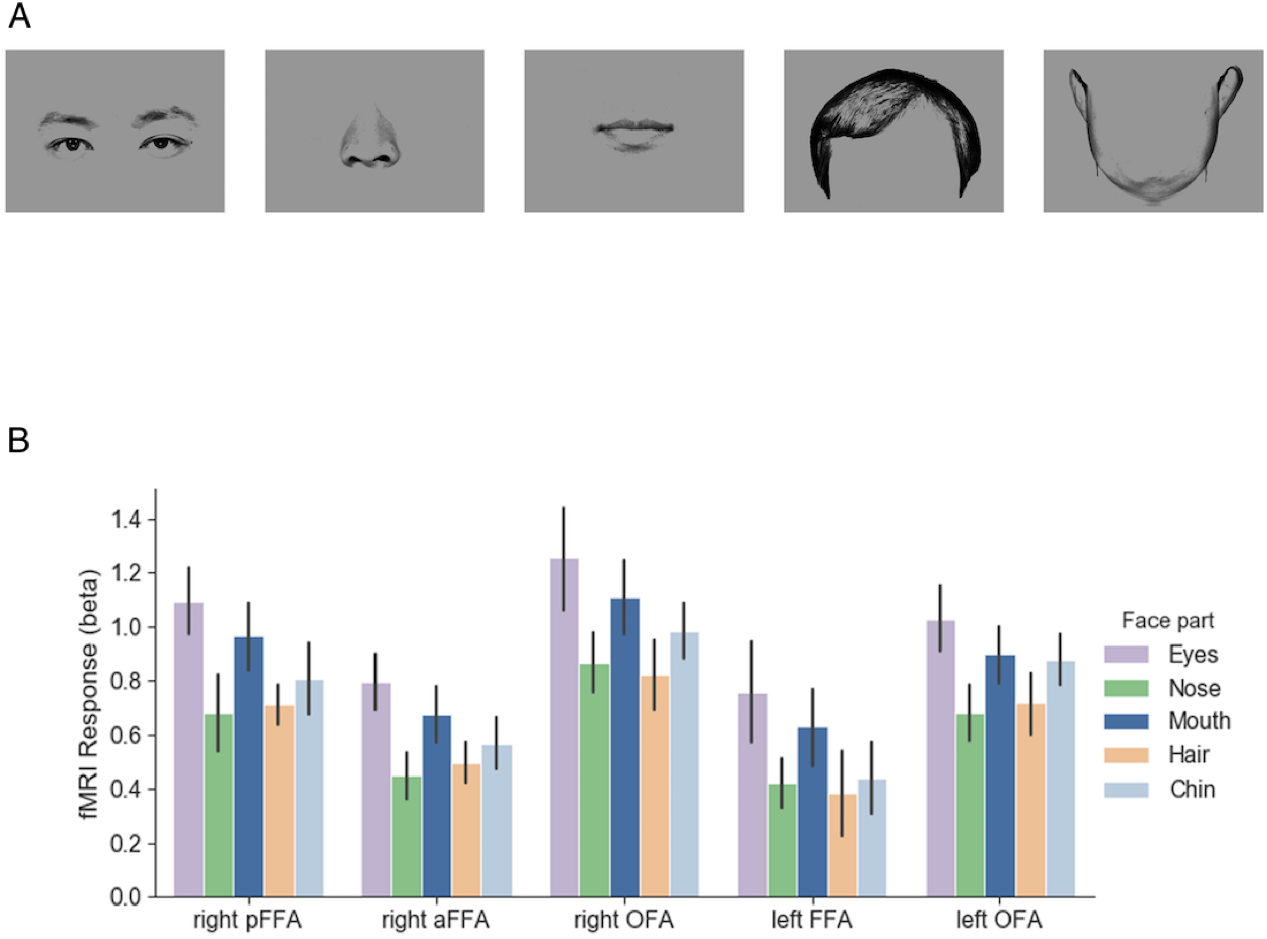
Stimuli and fMRI response maps. (A) Exemplars of face part stimuli used in the main experiment. The face parts were generated from 20 male faces. Each stimulus was presented around the fixation and participants performed a one-back task during the scan. (B) Average fMRI responses to different face parts in each face-selective region. Generally, eyes elicited higher responses than responses to nose, hair, and chin in most of the regions. No significant difference was observed between eyes and mouth responses. Error bars reflect ±1 SEM.

Considering that eyes and mouth are two dominant features in face perception (22, 23), and their response amplitudes were similar in face-selective regions, in the initial step, we compared the spatial patterns of neural responses to eyes with that to mouth within each ROI. Each pattern was first normalized to remove any overall amplitude difference between conditions. Then we directly contrasted the two patterns and projected the difference onto the inflated brain surface. A spatial pattern was observed in the right pFFA consistently across all participants (Figure 2). In the dimension parallel to the mid-fusiform gyrus, the posterior portion of the right pFFA was biased to respond more to eyes, whereas the anterior portion was biased to respond more to mouth. Note that in participant S2, the direction of MFS was more lateral-medial near the position of the right pFFA, and interestingly the eyes-mouth contrast map was oriented in the same direction, even though S2’s map may initially appear oriented differently from that of other participants. It suggests the anatomical orientation of MFS is highly correlated with such functional spatial organization of face parts.

**Figure 2.**
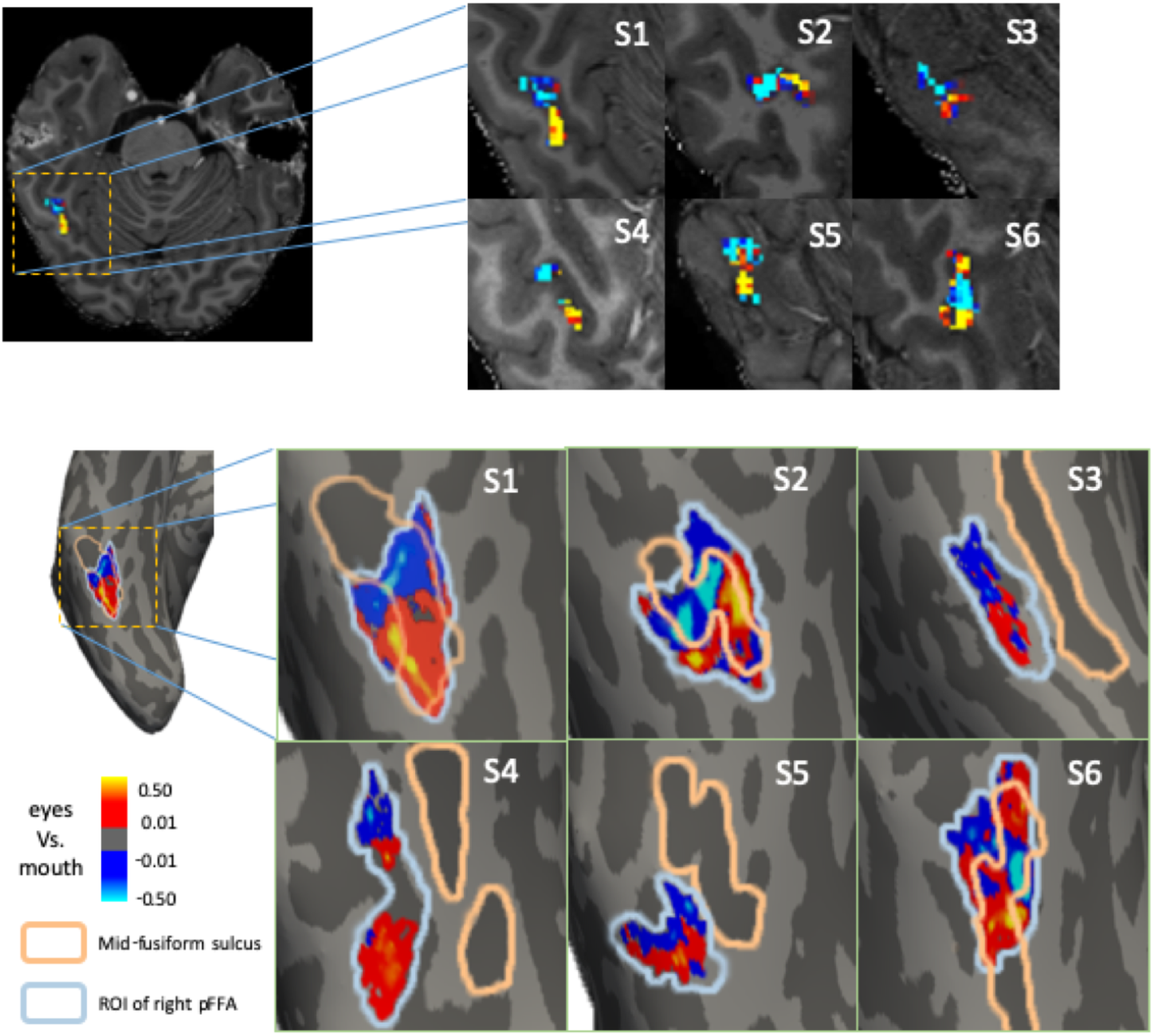
Contrast maps between normalized fMRI responses to eyes and mouth in the right pFFA illustrated in the volume (upper) or on inflated cortical surface (lower) of each participant. On the surface, the mid-fusiform sulcus is shown in dark gray with orange outline. The blue line outlines the right pFFA identified with an independent localizer scan. Aligned with the direction of mid-fusiform sulcus, the posterior part of right pFFA shows response bias to eyes (warm colors), while the anterior part illustrates mouth bias (cool colors). The posterior to anterior pattern is generally consistent across participants.

To further demonstrate such relationship, and also to provide a quantitative description of the spatial organization of face parts within right pFFA, we grouped voxels based on their location along the direction parallel to the MFS, and averaged the voxel responses at each location to generate the response profile on this posterior-anterior dimension (Figure 3A, see details in Method). The group-averaged results clearly showed that the difference between eyes and mouth signals consistently changed along the posterior-anterior direction in the right pFFA (Figure 3B). To quantify this trend, we further calculated the correlation coefficient between the eyes-mouth neural response differences and the position index along the posterior-anterior dimension (i.e., more posterior location was assigned with smaller value) in each participant. The group result revealed a significant negative correlation (t(5)=8.36, p=0.0004, Cohen’s d=3.41), confirming the consistency across participants that the posterior part of right pFFA was biased to eyes and anterior part was biased to mouth.

**Figure 3.**
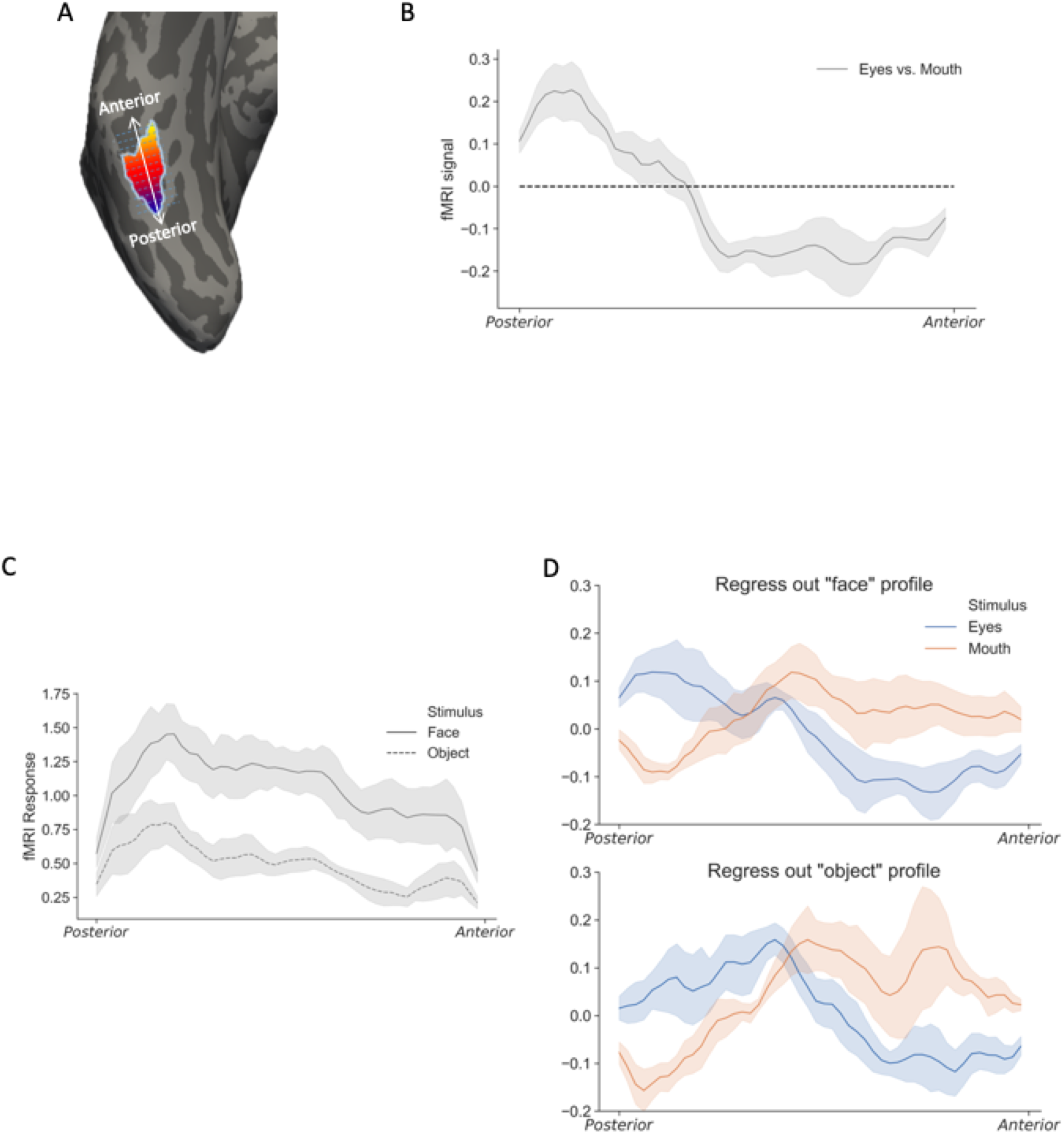
The spatial profiles of eyes and mouth responses along the posterior-anterior dimension. (A) To obtain the one-dimensional spatial profile of fMRI responses, a line was drawn parallel to the direction of mid-fusiform sulcus. Response from each vertex in the right pFFA was projected to the closest point on the line and averaged to generate the response profile. (B) The response profile of eyes vs mouth on the anterior-posterior dimension in right pFFA. The shaded regions reflect ±1 SEM. (C) The spatial profiles of whole faces and everyday objects in the right pFFA. Both profiles showed similar patterns, though the whole face responses were generally higher than object responses. (D) The spatial profile of individual face part responses, after regressing out the general fMRI response patterns elicited by either the whole faces (upper) or everyday objects (lower). In both cases, distinct spatial profiles were observed between eyes and mouth in the right pFFA.

The contrast map highlighted the differences between eyes and mouth responses. However, the original response patterns elicited by eyes and mouth share the same underlying general “face-related” pattern, which was subtracted out when contrasting the two response patterns. To extract the response profile of individual face parts, we used independently obtained response patterns of whole faces as the general face-related pattern and regressed it out from the eyes and mouth response patterns. With the general pattern regressed out, we observed distinct spatial profiles elicited by eyes and mouth in the right pFFA (Figure 3D top panel). The eye-biased voxels were more posterior than that of mouth-biased voxels, which is consistent with the contrast map shown in Figure 2.

Removing the general pattern helped to reveal the pattern of voxel biases for individual face parts. While removing the face-related general pattern achieved this goal, it is possible that removing the general face-related pattern distorted the parts generated response patterns since they share high-level visual information (i.e., face and eyes stimuli are both face-related). Therefore, it is important to check whether the parts specific patterns could be seen with removal of a common face-independent signal distribution. In five of the six participants, data were also obtained when they viewed everyday objects. Indeed, non-face objects generated significantly lower but spatially similar patterns of activation compared with faces across the right pFFA (Figure 3C). This result suggests that there is a general intrinsic BOLD sensitivity profile in the pFFA regardless of the stimuli. Thus we proceeded to use the response patterns of either faces or non-face everyday objects to regress out the intrinsic baseline profile from eyes and mouth response patterns, and plotted face part specific patterns along the posterior-anterior dimension. Consistently, results with object patterns removed showed clear posterior bias for eyes and anterior bias for mouth in the right pFFA (Figure 3D bottom panel).

To control for the potential contribution from retinotopic bias of the different face part conditions, in our experiment, all stimuli were presented at the fixation with a 1.3° horizontal jitter either to the left or to the right alternatively in different trials within a block. Even though the stimuli were centered on the fixation, because of the nature of the face parts (e.g., two eyes are apart, chin depicts the outline of the face), there were still small degrees (less than 3°) of retinotopic differences between the eyes and mouth conditions. To further rule out the retinotopic contribution, as well as to replicate our finding, we did two control experiments. In the first control experiment (Control Experiment 1), data were obtained with a single eye or mouth presented at either the near central (1.3°) or near peripheral (3.1°) location during the scan (see Figure S1A). This 2×2 (face parts x location) design allowed us to contrasted fMRI response patterns between face parts (single eye vs. mouth) regardless the stimulus location, or between locations (near central vs. near peripheral) regardless the face parts presented. Data from six participants were collected in the Control Experiment 1 and two of them (S1 and S5) also participated main experiment. In all participants, the eye vs. mouth contrast revealed spatial patterns in the right pFFA very similar to that in the main experiment (Figure S1B). However, contrasting fMRI responses between the near central and near peripheral location regardless the face parts failed to reveal consistent patterns across participants (Figure S1C). These results further support that the different fMRI response patterns we observed in the right pFFA were contributed by face feature differences rather than retinotopic bias. In the second control experiment (Control Experiment 2), we used top and bottom parts of the face as stimuli and counterbalance the stimulus location to verify the spatial organization in the right pFFA. With a 2×2 design (eyes vs. nose & mouth x present above vs. below fixation) (Figure S2A), consistent anterior-posterior spatial patterns in the right pFFA were observed in 8 participants (Figure S2B), which further corroborated our main finding of spatially organized representation of face parts in the right pFFA.

In addition to the two control experiments, we also measured the population receptive field (pRF) of each voxel in the right pFFA in three participants from the main experiment (Figure S3A) following established procedures (24–26). For each voxel, parameter x and y were calculated along with other parameters to represent the receptive field center on the horizontal (x) and vertical (y) axis in the visual field. Although generally more voxels in the right pFFA were bias to left visual field, which is consistent with previous report (27, 28), we observed no consistent spatial pattern in either x or y map of the right pFFA across participants (Figure S3B).

To examine the spatial patterns of response from eyes and mouth in other face-selectivity regions, similar analyses as in pFFA were applied to the fMRI response patterns in the right OFA, right aFFA, left FFA, and left OFA. For the left and right OFA, the posterior-anterior dimension was defined as the direction parallel to the occipitotemporal sulcus (OTS), where the OFAs were located in most participants. Among these regions, the right OFA also had distinct response patterns for eyes and mouth along the posterior-anterior dimension (Figure 4), similar to what we observed in the right pFFA. Group negative correlation was observed between the eyes-mouth differences and the posterior-anterior location of the right OFA (t(5) = 3.63, p=0.015, Cohen’s d=1.48). Such pattern was also observed in the Control Experiments. While the right OFA and right pFFA have been considered as sensitive to facial components and whole faces respectively, in our data they showed similar spatial profiles of eyes and mouth responses along the posterior-anterior dimension. This is consistent with, but adds some constraints to, the idea that the right pFFA may receive face feature information from right OFA for further processing (29, 30). In other face-selective regions, no consistent pattern was observed, as the correlations between the eyes-mouth difference and posterior-anterior location were not significant (ts<1.09, ps>0.32, see Figure 4A).

**Figure 4.**
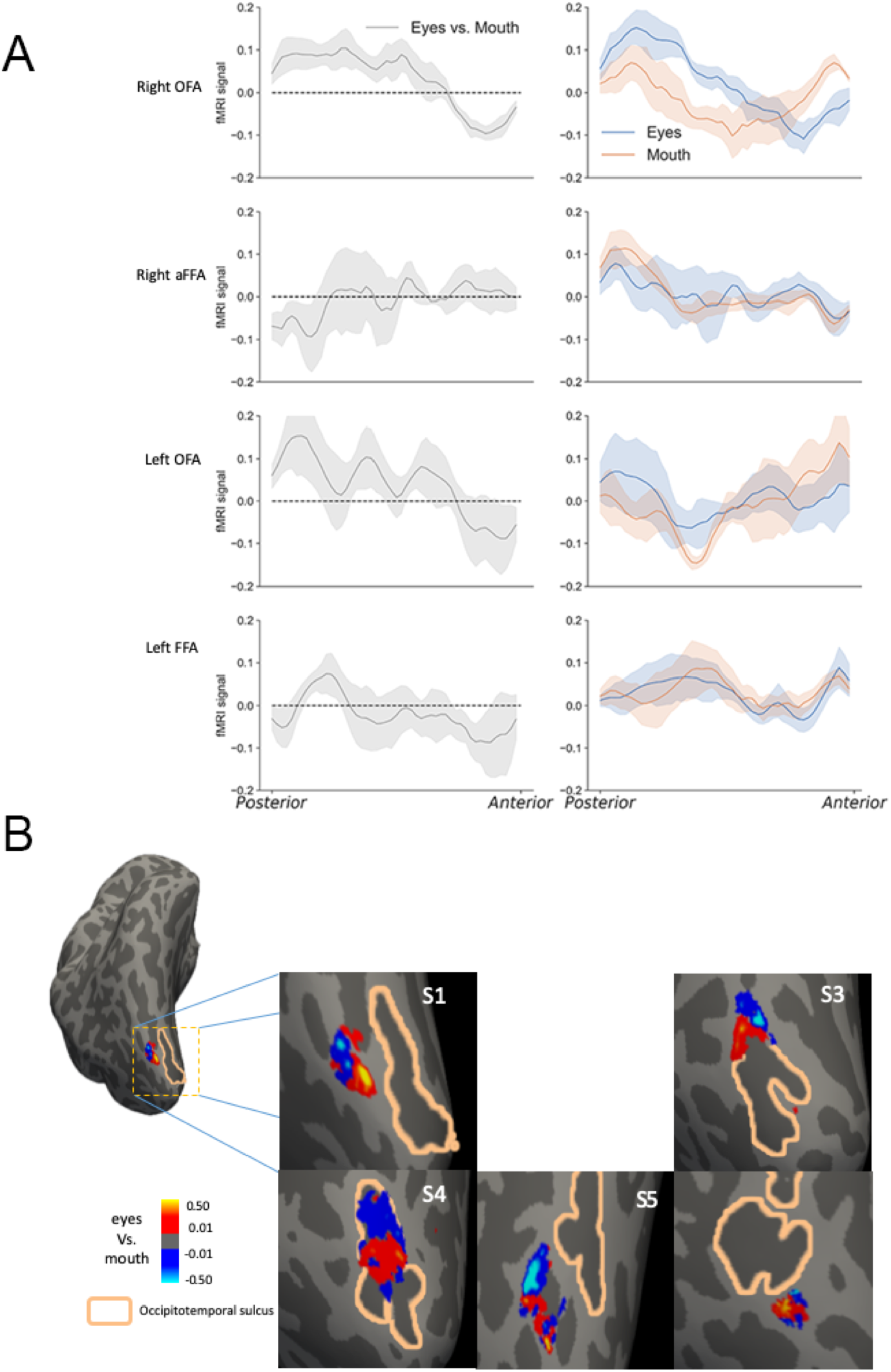
The spatial organizations of face parts in other face-selective regions. (A) The anterior-posterior neural response profiles of eyes and mouth in other face-selective regions. The contrast between normalized eyes and mouth response patterns in different regions are shown in the left column. The right column plots show the eyes and mouth response profiles with general response patterns regressed out. The right OFA (top panel) demonstrates different response profiles for eyes and mouth, similar to the observation in right pFFA. The shaded regions reflect ±1 SEM. (B) The eyes vs mouth contrast maps in right OFA in the main experiment. A consistent anterior-posterior pattern could be observed in each participant. Right OFA could not be identified in one of the six participants.

Beside the anterior-posterior dimension, the spatial representation of parts could organize in other spatial dimensions, such as the lateral-medial dimension in the VTC, or even in more complex nonlinear patterns. However, since the right pFFA located within the sulcus (MFS) in most of our participants, such that voxels distant from each other on the surface along the lateral-medial dimension could be spatially adjacent in the volume space, making it difficult to accurately reconstruct the spatial pattern along the lateral-medial dimension within the sulcus. Nevertheless, the finding of anterior-posterior organization of face parts is sufficient to demonstrate the existence of fine-scale feature map within object-selective regions.

Our stimuli also included nose, hair, and chin images, thus gave us a chance to examine their spatial profiles in each face-selective ROI as we did for eyes and mouth, though their neural responses were generally lower than that from eyes and mouth. Chin and mouth elicited similar response patterns along the anterior-posterior dimension in the right pFFA and right OFA after regressing out general spatial patterns (Figure 5A). By directly contrasting fMRI response patterns between eyes and chin, similar spatial profiles were revealed in the right pFFA and right OFA that the posterior part was biased to eyes and anterior part was biased to chin (ts>5.30, ps<0.01, see Figure 5B). We also observed a similar though less obvious profile in the left FFA (t(5)=2.68, p=0.04, Cohen’s d=1.09), but not in the right aFFA or left OFA (ts<0.41, ps>0.71).

**Figure 5.**
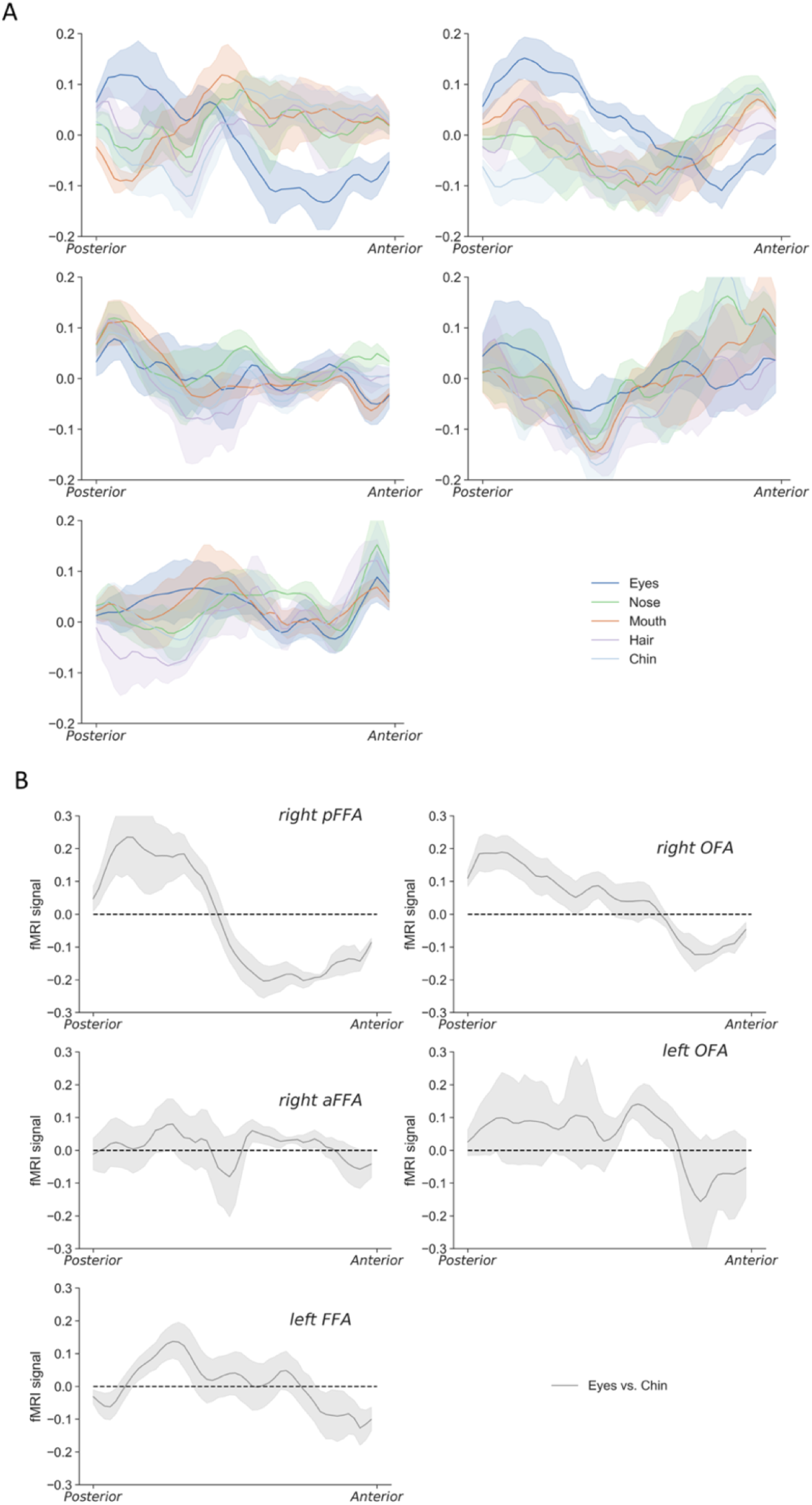
The anterior-posterior neural response profiles of all face parts (i.e., eyes, nose, mouth, hair, and chin) in five face-selective regions. (A) With general pattern regressed out, the chin showed similar response profiles as mouth in right pFFA and right OFA. (B) Contrasting normalized eyes and chin response patterns revealed consistent changes along the posterior-anterior dimension in right pFFA and right OFA. The shaded regions reflect ±1 SEM.

## Discussion

Our results reveal that within certain face-selective regions in the human occipito-temporal cortex, the neural representations of different facial features have consistent spatial organizations. Such fine-scale organizations are found similarly in the right pFFA and right OFA, but not in the more anterior right aFFA nor in the left hemisphere’s face-selective regions. In other words, fine-scale organizations for face parts exist at the early to intermediate stages of face processing hierarchy in the right hemisphere.

In the current study, five face parts (i.e., eyes, nose, mouth, hair, and chin) were tested, with eyes and mouth showed most distinct spatial organizations in the right pFFA and right OFA. No obvious spatial pattern was observed for nose and hair in face-selective regions, but it would be premature to conclude that there is no fine-scale spatial organization for their neural representations. For one, the nose and hair stimuli elicited lower fMRI responses compared with eyes and mouth stimuli, making it more difficult to detect potential spatial patterns. The observation that eyes and mouth elicited most differential patterns is consistent with them providing more information about faces than other features in face processing (22, 23). The dominance of eyes and mouth in face-selective regions could be considered as a form of cortical magnification of more informative features, a common principle of functional organization in sensory cortex (31–33).

The discovery that some face parts are represented within the face-processing regions with fine-scale spatial organization improve our understanding about how functional representations are physically applied to anatomical structures in the VTC. To further explore the neural models about object processing in the VTC, it is important to ask what kinds of constrain, functional or anatomical, result in such fine-scale organization? Many of the visual cortical areas have retinotopic maps, indeed having a retinotopic representation of the visual world is one of the key ways to define a visual cortical area. Along occipitotemporal processing stream, visual areas increasingly become more specialized in processing certain features and object categories. What is the relationship between a potential spatial organization of face part representations and the spatial relationship of face parts in a real face?

A recent study has revealed that in the inferior occipital gyrus, where the OFA located, both tunings for retinotopic location and face parts (34). Although the tuning peak maps were idiosyncratic across individuals, the two tuning maps were correlated within individuals, suggesting a relationship between face parts configuration and their typical retinotopic configuration. Our findings provide additional support for face part turning in the OFA, and further reveal that there exists a consistent organization of face part tuning *across* individuals. More importantly, our finding of spatial organization of face part tuning in the pFFA indicates that although the organization of feature tuning could be constrained by the retinotopic tuning in occipital cortex, a more abstract feature tuning could still be spatially organized in cortical areas with absent or minimal retinotopic property in the later stages of VTC.

Another previous study also tested the idea of “faciotopy”, that there are cortical patches representing different face features within a face-selective region and the spatial organization of these feature patches on the cortical surface would reflect the physical relationships of face features (11). Their results showed that in the OFA and FFA, the differences between neural response patterns of face parts were correlated with physical distances between face parts in a face. Our results support the existence of stable organization of face features in the right OFA and right pFFA, especially for eyes and mouth. The possible mechanism underlying such faciotopy organization is the local-to-global computation, that physically adjacent face parts interact more than parts far apart from each other during the early stages of processing, thus it is more efficient for them to have neural representations near each other. However, in the current study, we did not find the posterior bias pattern for hair as we did for eyes, even though hair and eyes are spatially adjacent. which could be caused by the hair being generally less invariant and less informative in the face identification.

Another potential explanation could be that while both contribute to face identification, eyes and mouth are differentially involved in different neural networks and have distinct functional connectivity profiles with other brain regions. Specifically, the mouth region provides more information for language processing and audio-visual communication perception, thus it may be more connected to the language processing system. Meanwhile, the eyes are more important in face detection and eye gaze signifies interest, thus it may be more connected to the attention system. Previous studies have already found the connectivity profiles could predict the functional selectivity in the VTC, thus it would be interesting to examine whether the face part organization in the pFFA could be predicted using functional or anatomical connectivity to the other brain regions in the future studies.

Among five face-selective regions we examined, only the right pFFA and right OFA exhibited distinct fMRI response patterns for eyes and mouth. In the face processing network, face parts are believed to be represented in the posterior regions such as the OFA (29, 35, 36). Part information is transmitted to anterior regions to be further integrated to form holistic face representations (10, 37). In that sense, the more anterior regions in the face processing network are more responsible for representing integrated face information such as gender or identity rather than individual face parts (38, 39). Consistent with this idea, at the right aFFA, there is no obvious spatial organization of face parts. Meanwhile, a clear hemispheric difference was found in our results that the distinct spatial response patterns for face parts were observed in the right but not the left hemisphere, which is consistent with previous findings that compared with left FFA, right FFA is more sensitive to face specific features (40) and configural information (41). The right face selective regions may carry more distinct face feature representation than left hemisphere as face-processing network is more dominant in right hemisphere.

Much progress has been made in our understanding of object feature representation in the VTC during the past decade, especially with the view of feature space representation (3, 8). Consequently, we now believe that a large number of features are represented for object recognition, but how does our brain physically organize such complex feature representations in the VTC? One possible solution is that these feature representations manifest in different spatial scales. For more general features the representation manifests at large spatial scale across the whole VTC (e.g., large/small, animate/inanimate), and for more specific features such as face parts, it manifests at finer spatial scales within specific object processing regions. Under this view, we would expect more fine-scale feature organizations to be revealed with more advanced neural imaging tools, which are critical for fully understanding the neural algorithm of object processing in the VTC.

## Materials and Methods

### Participants

Six (3 females) human participants were recruited in the main experiment. Six (5 females) participants (two of them also participated main experiment) were recruited in the Control Experiment 1. Three participants (2 females) from main experiment finished the pRF experiment. Ten participants (1 female) were recuited in the Control Experiment 2, but in two participants right pFFA was failed to be localized, thus we excluded these two participants from the analyses. All participants were between the ages of 21 and 27, right-handed, and had normal or corrected to normal visual acuity. They were recruited from the Chinese Academy of Sciences community with informed consent and received payment for their participation in the experiment. The experiment was approved by the Committee on the Use of Human Subjects at the Institute of Biophysics of Chinese Academy of Sciences.

### Stimuli and Experimental design

In the main experiment, for face stimuli, 20 unique front-view Asian male face images were used. Each face image was gray-scaled and further divided into five parts (i.e., eyes, nose, mouth, hair, and chin. See Figure 1A). Twenty unique gray-scaled everyday objects were used as comparison stimuli. The full face and object images on average subtended around 5° x 7°. For stimuli used in localizer scans, video clips of faces, objects, and scrambled objects were used (For detail see (42)).

There were total of seven stimulus conditions (i.e., eyes, nose, mouth, hair, chin, whole face, and object condition). Each main experimental run contained two blocks of each stimulus condition. In the scan of participant S6, the object condition was not included. Each stimulus block lasted 16 sec and contained 20 images of the same type. Each image was presented for 600 msec at fixation and followed by a 200-msec blank interval. There was a 16-sec blank fixation block at the beginning, the middle, and the end of each run. Participants performed a 1-back task that they were asked to press a button when two successive images were the same. To balance the spatial property in the visual field of different images, each image was presented at a slightly shifted location, 1.3° either to the left or to the right of the fixation alternately in different trials within a block. Participants were instructed to maintain central fixation throughout the task.

Each localizer run contained four 18-sec blocks of each of the three stimulus conditions (i.e., faces, everyday non-face objects, and scrambled objects) shown in a balanced block order. The 12 stimulus blocks were interleaved by three 18-sec fixation blocks inserted at the beginning, middle and end of each run. Each block contained 6 video clips of a given stimulus category, each presented for 3 sec. Participants were asked to watch the video without any task. No fixation point was presented during the scan.

The 8 experimental runs and the 2 localizer runs were completed within the same scan session for each participant.

In the Control Experiment 1, we used a similar block design as that in the main experiment. There were six kinds of stimulus blocks (single eye near central, single eye near peripheral, mouth near central, mouth near peripheral, whole face, object) and each of them repeated three times in a single run. Each participant completed four runs and two localizer runs. In the eye near central condition, single left eye images were presented at 1.3° either to the left or to the right of the fixation alternately in different trials within a block. In the eye near peripheral condition, single left eye images were presented at 3.1° either to the left or to the right of the fixation. The central and peripheral locations were chosen to match the locations of eyes and mouth in the main experiment. Stimuli in mouth near central and mouth near peripheral conditions were presented in the same locations as in two eye conditions respectively. Whole face and object conditions were the same as in the main experiment.

In the pRF experiment, we adopted stimuli and analysis code from analyzePRF package (http://kendrickkay.net/analyzePRF/). There were total of four conditions (i.e., clockwise wedges, counterclockwise wedges, expanding rings, contracting rings). The angular span of the wedges was 45°, and it revolved for 32 seconds per cycle. In the rings conditions, the rings swept 28 seconds per cycle with 4 seconds of rest followed. Colored object images were presented on the wedges or rings. The rings and wedges were presented within a radius of 10°. For each run there was a 22-sec blank fixation block at the beginning and the end. Participants performed a change detection task that they pressed a button whenever the fixation color changed. In each run, only one kind of PRF stimulus was presented and repeated 8 cycles. Each participant finished 4 different pRF runs.

In the Control Experiment 2, similar block-design as in main experiment was used. Four face part conditions (top vs bottom part x present location) were included in the experiment (Figure S2A). The top part contained eyes (4.02° x 12.08°) and the bottom part contained nose and mouth (8.08° x 12.08°). To engage observers’ attention on the stimuli, a randomly selected four images in each block moved slightly either to the left or right during stimulus presentation. Observers were asked to judge the directions of these movements. Same localizer runs as in the main experiment were included for each participant.

### FMRI scanning

MRI data were collected on Siemens Magnetom 7 Tesla MRI system (passively shielded, 45mT/s slew rate) (Siemens, Erlangen, Germany), with a 32-channel receive 1-channel transmit head coil (NOVA Medical, Inc, Wilmington, MA, USA), at the Beijing MRI Center for Brain Research (BMCBR). High-resolution T1-weighted anatomical images (0.7 mm isotropic voxel size) were acquired with a MPRAGE sequence (256 sagittal slices, acquisition matrix = 320 × 320, Field of view = 223 × 223 mm, GRAPPA factor = 3, TR = 3100 ms, TE = 3.56 ms, TI = 1250ms, flip angle = 5°, pixel bandwidth = 182 Hz per pixel). Proton density (PD)-weighted images were acquired with same voxel size and FOV (256 sagittal slices, acquisition matrix = 320 × 320, Field of view = 223 × 223 mm, GRAPPA factor = 3, TR = 2340 ms, TE = 3.56 ms, flip angle = 5°, pixel bandwidth = 182 Hz). GE-EPI sequences was used to collect functional data in the main experiment (TR = 2000ms, TE = 18ms, 1.2mm isotropic voxels, FOV = 168 × 168 mm, image matrix = 140 × 140, GRAPPA factor = 3, partial Fourier 6/8, 31 slices of 1.2mm thinkness, flip angle is about 80, pixel bandwidth = 1276 Hz per pixel). During the scan, GE-EPI images with reversed phase encoding direction from experiment functional scan were collected to correct the spatial distortion of EPI images(43). Dielectric pads were placed on both sides of the head to improve B1 efficiency in the temporal cortex(44), while bite-bar was used to reduce head movements for each participant. During the functional scan, respiration and pulse signals were recorded simultaneously. GE-EPI sequences with same resolution as in the main experiment was used in the control and pRF experiment (TR = 2000ms, TE = 22ms, 1.2mm isotropic voxels, FOV = 180 × 180 mm, image matrix = 150 × 150, GRAPPA factor = 2, partial Fourier 6/8, 31 slices of 1.2mm thinkness, flip angle is about 80, pixel bandwidth = 1587 Hz per pixel). Dielectric pads were placed on the right side of the head.

### Data analysis

Anatomical data were analyzed with FreeSurfer (Cortechs Inc, Charlestown, MA) and custom MATLAB codes. To enhance the contrast between white and gray matter, T1-weighted images were divided by PD-weighted images (45). Anatomical data were further processed with FreeSurfer to reconstruct the cortical surface models.

Functional data were analyzed with AFNI (http://afni.nimh.nih.gov), FreeSurfer, fROI (http://froi.sourceforge.net)), and custom MATLAB codes. Data preprocessing included slice-timing correction, motion correction, removing physiological noise with respiration and pulse signals, distortion correction with reversed phase encoding EPI images, and intensity normalization. For the localizer runs only, spatial smoothing was applied (Gaussian kernel, 2 mm full width at half maximum). After preprocessing, function images were co-registered to anatomic images for each participant. To obtain the average response amplitude for each voxel in the specific stimulus condition for each individual observer, voxel time courses were fitted by a general linear model (GLM), whereby each condition was modeled by a boxcar regressor (matched in stimulus duration) and then convolved with a gamma function (delta = 2.25, tau = 1.25). The resulting beta weights were used to characterize the averaged response amplitudes.

The face-selective ROIs were identified by contrasting functional data between face and everyday-object conditions in the localizer runs. Specifically, FFA and OFA was defined as the set of continuous voxels in fusiform gyrus and inferior occipital gyrus, respectively, that showed significantly higher response to faces than to objects (p < 0.01, uncorrected). We were able to identify right pFFA, right anterior FFA (right aFFA), right OFA, and left FFA in all six participants. The left OFA were successfully identified in five out of six participants. In each ROI, to remove the vein signal in the functional data, voxels of which signal changes to face stimuli were larger than 4% were excluded in further analysis.

For the main experimental data, to remove the general fMRI response pattern shared among different face parts, response patterns from whole faces or everyday objects were regressed out from response patterns of each individual face part. Whole face or object response in each voxel was used to predict the individual part response with linear regression algorithm, and the residuals across voxels were considered as the individual part response pattern with general response pattern removed. To extract the trend of the fMRI response pattern along anterior-posterior dimension in the FFA, we first drew a line along the mid-fusiform sulcus on the cortical surface of each participant. For all vertices within the FFA ROI, we calculated their shortest (orthogonal) distances to the line, and projected the neural response of all voxels in the FFA ROI to the line along the mid-fusiform sulcus, and obtained the averaged response on each point along the line to get the response profiles (see Figure 3A). Similar analysis was done for OFA with the line drawn along the inferior occipital sulcus.

For the control experiments, same data processing steps as in the main experiment were applied to extract the spatial patterns of different conditions. For the pRF data, fMRI respond time course of each voxel was fit with compressive spatial summation (CSS) model (http://kendrickkay.net/analyzePRF/). To determine the center location (x, y) of each voxel’s population receptive field, CSS used an isotropic 2D Gaussian and a static power-low nonlinearity to model the fMRI response. In each voxel, model fitness can be quantified as the coefficient of determination between model and data (R^2^). We only included the pRF results of voxels with R^2^ higher than 2%.

## Data availability

Source data for the results are available on Dryad (https://datadryad.org/stash/share/jZx0MR_ueTu0pwAixlvFAHfQetoGcW28sdXAyJ5_2Uk).

## Acknowledgments

This work was supported by funds from CAS Pioneer Hundred Talents Program, NSFC grant (No. 31800966), Strategy Priority Research Program of Chinese Academy of Science (No. XDB32020200), and Beijing Science and Technology Project (No. Z181100001518002). The authors would like to thank Dr. Chencan Qian and Dr. Zihao Zhang for their help during data collection and analysis.

## Author Contributions

J.Z., P.Z., and S.H. designed the experiments; J.Z., Y.J., Y.S., and P.Z. conducted the experiments; J.Z., Y.J., and Y.S. analyzed the data; J.Z., P.Z., and S.H. wrote the paper.

## Declaration of Interests

The authors declare no competing interests.

---

**Figure S1.**
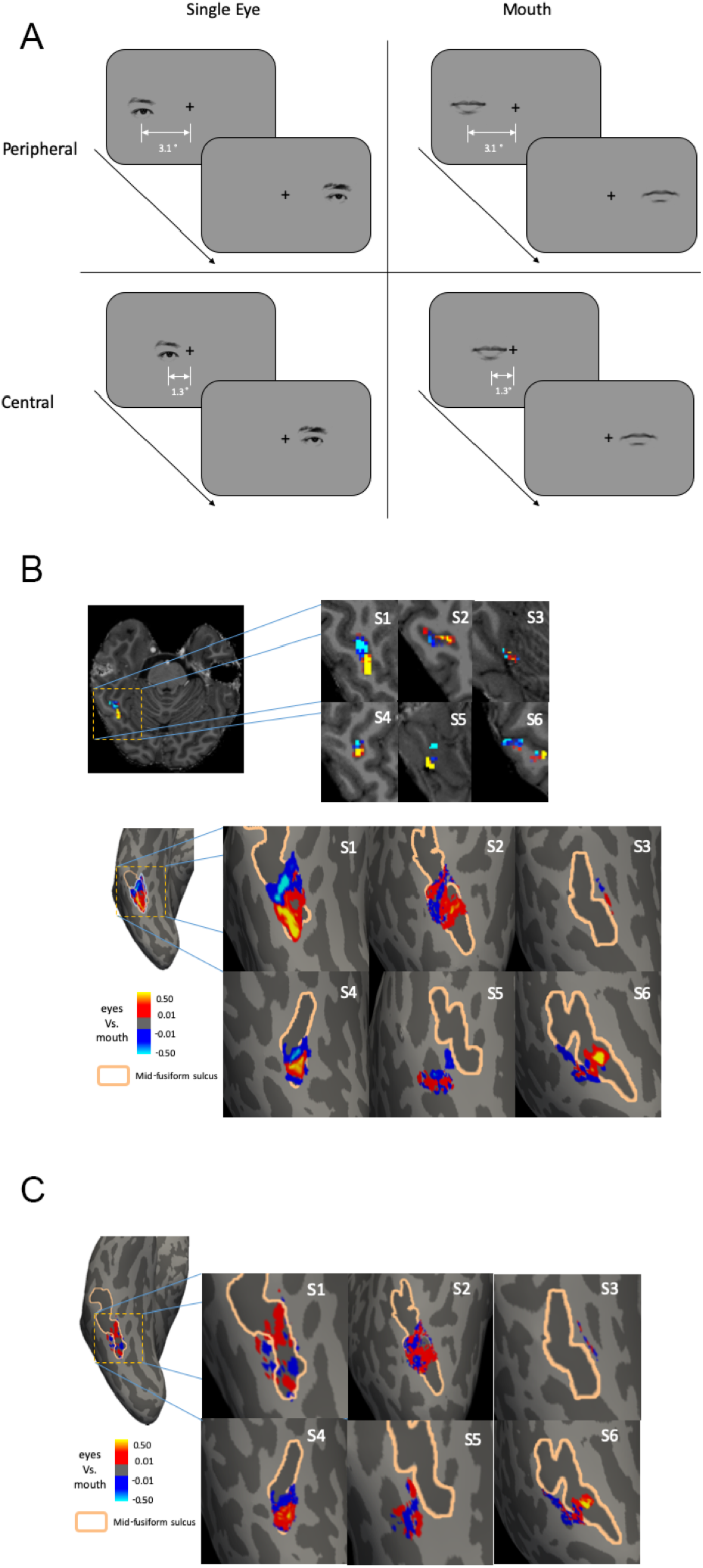
Stimuli and results of Control Experiment 1. (A) Four conditions (i.e., single eye near peripheral, single eye near central, mouth near peripheral, mouth near central) were include in the Control Experiment 1. For the near peripheral conditions, eye or mouth were presented at 3.1° either to the left or to the right of the fixation. For the near central conditions, face parts were presented at 1.3° left or right to the fixation. (B) The contrast maps between eyes and mouth regardless of location in each participant, showing clear anterior-posterior trend. (C) The contrast maps of near peripheral vs. near central location regardless of face parts, no consistent pattern is observed.

**Figure S2.**
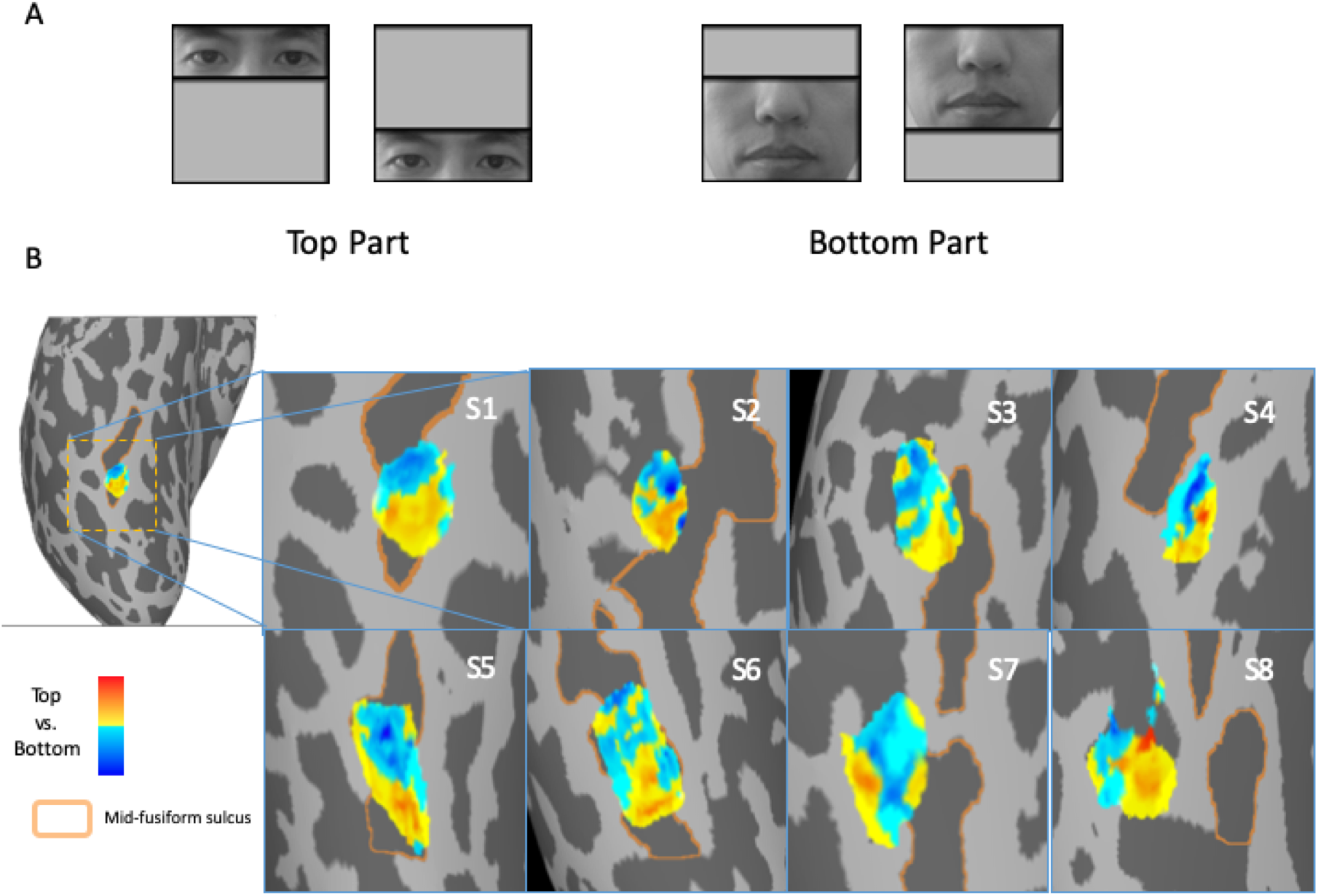
(A) Stimulus examples from Control Experiment 2 that top or bottom face parts presented at upper or lower visual field. (B) The anterior-posterior neural response profiles of face parts in right pFFA in eight participants.

**Figure S3.**
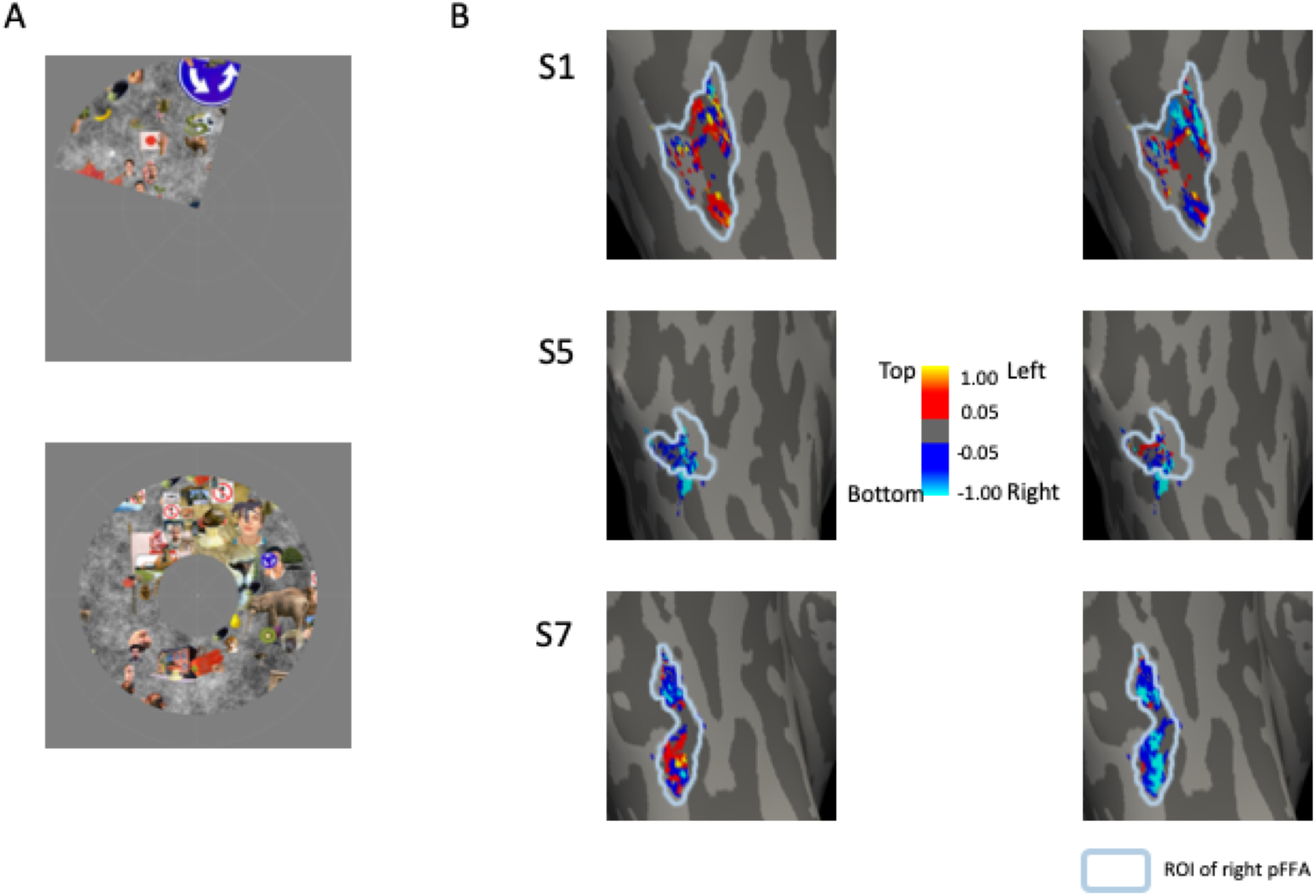
Stimuli and results of pRF experiment. (A) Wedge and ring stimuli used in the pRF experiment. (B) Visualization of receptive field center of each voxel in the right pFFA outlined in blue. The maps represent the locations of receptive field center on the horizontal (left column) or vertical (right column) axis. No consistent spatial pattern is found across different participants.

## Notes

### Competing Interest Statement

The authors have declared no competing interest.

## References

1. U. Hasson, M. Harel, I. Levy, R. Malach, Large-scale mirror-symmetry organization of human occipito-temporal object areas. Neuron 37, 1027–41 (2003).

2. M. Spiridon, B. Fischl, N. Kanwisher, Location and spatial profile of category-specific regions in human extrastriate cortex. Hum. Brain Mapp. 27, 77–89 (2006).

3. P. Bao, L. She, M. McGill, D. Y. Tsao, A map of object space in primate inferotemporal cortex. Nature 583, 103–108 (2020).

4. G. G. Blasdel, G. Salama, Voltage-sensitive dyes reveal a modular organization in monkey striate cortex. Nature 321, 579–585 (1986).

5. T. Bonhoeffer, A. Grinvald, lso-orientation domains in cat visual cortex are arranged in pinwheel-like patterns. 353, 3 (1991).

6. D. H. Hubel, T. N. Wiesel, Ferrier lecture. Functional architecture of macaque monkey visual cortex. Proc. R. Soc. Lond. B Biol. Sci. 198, 1–59 (1977).

7. M. Weliky, W. H. Bosking, D. Fitzpatrick, A systematic map of direction preference in primary visual cortex. Nature 379, 725–728 (1996).

8. L. Chang, D. Y. Tsao, The Code for Facial Identity in the Primate Brain. Cell 169, 1013-1028.e14 (2017).

9. W. A. Freiwald, D. Y. Tsao, M. S. Livingstone, A face feature space in the macaque temporal lobe. Nat. Neurosci. 12, 1187–1196 (2009).

10. J. Zhang, J. Liu, Y. Xu, Neural Decoding Reveals Impaired Face Configural Processing in the Right Fusiform Face Area of Individuals with Developmental Prosopagnosia. J. Neurosci. 35, 1539–1548 (2015).

11. L. Henriksson, M. Mur, N. Kriegeskorte, Faciotopy—A face-feature map with face-like topology in the human occipital face area. Cortex 72, 156–167 (2015).

12. K. Cheng, R. A. Waggoner, K. Tanaka, Human Ocular Dominance Columns as Revealed by High-Field Functional Magnetic Resonance Imaging. Neuron 32, 359–374 (2001).

13. N. R. Goncalves, et al., 7 Tesla fMRI Reveals Systematic Functional Organization for Binocular Disparity in Dorsal Visual Cortex. J. Neurosci. 35, 3056–3072 (2015).

14. S. Nasr, J. R. Polimeni, R. B. H. Tootell, Interdigitated Color- and Disparity-Selective Columns within Human Visual Cortical Areas V2 and V3. J. Neurosci. 36, 1841–1857 (2016).

15. M. Schneider, V. G. Kemper, T. C. Emmerling, F. De Martino, R. Goebel, Columnar clusters in the human motion complex reflect consciously perceived motion axis. Proc. Natl. Acad. Sci. 116, 5096–5101 (2019).

16. E. Yacoub, N. Harel, K. Ugurbil, High-field fMRI unveils orientation columns in humans. Proc. Natl. Acad. Sci. 105, 10607–10612 (2008).

17. J. Zimmermann, et al., Mapping the Organization of Axis of Motion Selective Features in Human Area MT Using High-Field fMRI. PLoS ONE 6, e28716 (2011).

18. K. S. Weiner, et al., The mid-fusiform sulcus: A landmark identifying both cytoarchitectonic and functional divisions of human ventral temporal cortex. NeuroImage 84, 453–465 (2014).

19. J. Caspers, et al., Cytoarchitectonical analysis and probabilistic mapping of two extrastriate areas of the human posterior fusiform gyrus. Brain Struct. Funct. 218, 511–526 (2013).

20. K. Grill-Spector, K. S. Weiner, The functional architecture of the ventral temporal cortex and its role in categorization. Nat. Rev. Neurosci. 15, 536–548 (2014).

21. S. Lorenz, et al., Two New Cytoarchitectonic Areas on the Human Mid-Fusiform Gyrus. Cereb. Cortex, bhv225 (2015).

22. P. G. Schyns, L. Bonnar, F. Gosselin, Show Me the Features! Understanding Recognition From the Use of Visual Information. Psychol. Sci. 13, 402–409 (2002).

23. M. Wegrzyn, M. Vogt, B. Kireclioglu, J. Schneider, J. Kissler, Mapping the emotional face. How individual face parts contribute to successful emotion recognition. PLOS ONE 12, e0177239 (2017).

24. S. O. Dumoulin, B. A. Wandell, Population receptive field estimates in human visual cortex. NeuroImage 39, 647–660 (2008).

25. K. N. Kay, J. Winawer, A. Mezer, B. A. Wandell, Compressive spatial summation in human visual cortex. J. Neurophysiol. 110, 481–494 (2013).

26. N. Kriegeskorte, et al., Matching categorical object representations in inferior temporal cortex of man and monkey. Neuron 60, 1126–1141 (2008).

27. K. N. Kay, K. S. Weiner, K. Grill-Spector, Attention Reduces Spatial Uncertainty in Human Ventral Temporal Cortex. Curr. Biol. 25, 595–600 (2015).

28. D. F. Nichols, L. R. Betts, H. R. Wilson, Position selectivity in face-sensitive visual cortex to facial and nonfacial stimuli: an fMRI study. Brain Behav. 6 (2016).

29. J. Liu, A. Harris, N. Kanwisher, Perception of face parts and face configurations: an FMRI study. J. Cogn. Neurosci. 22, 203–211 (2010).

30. Q. Zhu, J. Zhang, Y. L. L. Luo, D. D. Dilks, J. Liu, Resting-State Neural Activity across Face-Selective Cortical Regions Is Behaviorally Relevant. J. Neurosci. 31, 10323–10330 (2011).

31. A. Cowey, E. T. Rolls, Human cortical magnification factor and its relation to visual acuity. Exp. Brain Res. 21, 447–454 (1974).

32. P. M. Daniel, D. Whitteridge, The representation of the visual field on the cerebral cortex in monkeys. J. Physiol. 159, 203–221 (1961).

33. W. Penfield, E. Boldrey, SOMATIC MOTOR AND SENSORY REPRESENTATION IN THE CEREBRAL CORTEX OF MAN AS STUDIED BY ELECTRICAL STIMULATION. Brain 60, 389–443 (1937).

34. B. de Haas, M. I. Sereno, D. S. Schwarzkopf, Inferior occipital gyrus is organised along common gradients of spatial and face-part selectivity. J. Neurosci. (2021) https://doi.org/10.1523/JNEUROSCI.2415-20.2021 (June 15, 2021).

35. L. R. Arcurio, J. M. Gold, T. W. James, The response of face-selective cortex with single face parts and part combinations. Neuropsychologia 50, 2454–2459 (2012).

36. D. Pitcher, V. Walsh, G. Yovel, B. Duchaine, TMS Evidence for the Involvement of the Right Occipital Face Area in Early Face Processing. Curr. Biol. 17, 1568–1573 (2007).

37. P. Rotshtein, R. N. Henson, A. Treves, J. Driver, R. J. Dolan, Morphing Marilyn into Maggie dissociates physical and identity face representations in the brain. Nat Neurosci 8, 107–13 (2005).

38. W. A. Freiwald, D. Y. Tsao, Functional Compartmentalization and Viewpoint Generalization Within the Macaque Face-Processing System. Science 330, 845–851 (2010).

39. S. M. Landi, W. A. Freiwald, Two areas for familiar face recognition in the primate brain. Science 357, 591–595 (2017).

40. M. Meng, T. Cherian, G. Singal, P. Sinha, Lateralization of face processing in the human brain. Proc. R. Soc. B Biol. Sci. 279, 2052–2061 (2012).

41. B. Rossion, et al., Hemispheric asymmetries for whole-based and part-based face processing in the human fusiform gyrus. J Cogn Neurosci 12, 793–802 (2000).

42. D. Pitcher, D. D. Dilks, R. R. Saxe, C. Triantafyllou, N. Kanwisher, Differential selectivity for dynamic versus static information in face-selective cortical regions. NeuroImage 56, 2356–2363 (2011).

43. P. S. Morgan, R. W. Bowtell, D. J. O. McIntyre, B. S. Worthington, Correction of spatial distortion in EPI due to inhomogeneous static magnetic fields using the reversed gradient method. J. Magn. Reson. Imaging 19, 499–507 (2004).

44. W. M. Teeuwisse, W. M. Brink, A. G. Webb, Quantitative assessment of the effects of high-permittivity pads in 7 Tesla MRI of the brain. Magn. Reson. Med. 67, 1285–1293 (2012).

45. P.-F. Van de Moortele, et al., T1 weighted brain images at 7 Tesla unbiased for Proton Density, T21 contrast and RF coil receive B1 sensitivity with simultaneous vessel visualization. NeuroImage 46, 432–446 (2009).

